# Teatime for *Triticum* – (how) can the presence of plants slow down decomposition?

**DOI:** 10.64898/2026.03.19.712830

**Authors:** Jennifer Michel, Alice Quenon, Mayliss Persyn, Anna Xayphrarath, Adrien Blum, Vincent Leemans, Da Cao, Sara Sanchez-Moreno, Hervé Vanderschuren, Dominique Van Der Straeten, Markus Weinmann, Jordi Moya-Laraño, Pierre Delaplace

## Abstract

Decomposition of organic matter is a key process in soils contributing to carbon and nutrient cycling. To identify management strategies for agroecosystems that reduce nutrient losses while maximizing plant growth, it is important to understand which parameters determine decomposition rates. This study therefore investigated how the presence of winter wheat (*Triticum aestivum* var. Asory) affects decomposition in a controlled Ecotron setup with two soil types with varying organic matter content across three simulated climates (2013, 2068, 2085). Using the tea bag index, interstitial soil pore water analyses, microbial biomass quantification, bacterial and fungal gene abundance, and soil respiration measurements, we tested the hypotheses that plant exudates would enhance decomposition rate and microbial biomass, while plant nitrogen uptake would deplete soil nitrate, potentially mitigated by fertilization. Contrary to expectations, decomposition rates were lower in planted than in unplanted soils, suggesting resource competition between plants and microbes. No significant differences were observed in microbial biomass or respiration due to plant presence, and fertilization effects on nitrate or microbial mineralization were undetectable, likely due to rapid turnover of organic molecules including uptake by plants and microbes. Mechanistically, fungi and soil humidity were more important for decomposition than bacteria or temperature. The findings corroborate climate impacts on decomposition but also indicate microbial resilience and highlight the potential of management strategies like cover crops, adjusted planting dates and crop residual management which can contribute to healthy soils by sustaining carbon and nutrient cycling.

## 1. Introduction

Decomposition is a fundamental ecosystem process that regulates carbon and nutrient cycling, influencing soil fertility and plant productivity in agricultural systems. In the context of climate change, understanding how biotic factors, such as the presence of crop plants, interact with abiotic conditions to modulate decomposition is essential for sustainable farming. Plants can potentially accelerate decomposition through root exudates, particularly carbohydrate-rich compounds that fuel microbial activity and mineralization^[1,2,3,4]^. Conversely, plants may compete with microbes for resources like nitrogen, slowing decomposition via nutrient limitation^[5,6,7,8]^. This duality is particularly relevant for crops like winter wheat (*Triticum aestivum*), where effective nutrient management may determine how yields evolve under future climate^[9,10]^.

Despite extensive research on decomposition, few studies have focused explicitly on the effect of presence or absence of living plants, despite the relevance for agricultural land management. The plant effect has been studied even less under predicted future meteorological conditions, a critical research gap as projected climate scenarios with warmer temperatures and variable precipitation could exacerbate or buffer nutrient losses, depending on microbial communities and soil organic matter (SOM) content, which affects microbial resilience and water retention^[11,12,13]^. Here, we test how winter wheat presence influences decomposition, microbial biomass, bacterial and fungal gene abundance, respiration, and glucose and nitrate dynamics in presence or absence of plants based on an Ecotron experiment. The Ecotron experiment exposed cropping systems to the meteorological conditions of three different years across a climate gradient representative of climate change in the 21^st^ century in Central Europe and used two contrasting soil types (low vs. high SOM). We hypothesized that plants enhance decomposition via exudates but deplete nitrate through uptake, with fertilization compensating for the latter. These insights aim to inform adaptive agricultural practices, emphasizing the role of plant-microbe interactions in maintaining soil health.

## 2. Material & methods

### 2.1 Hypotheses

This experiment focused on decomposition and nitrate availability in relation to the presence/absence of plants and in relation to nitrogen fertilization.

The first set of hypotheses was based on the premise that plants exude carbohydrate-rich substances which increase microbial mineralization^[1,2]^. From this we deduced that

HP1a: Decomposition (tea bag index) is higher in planted than in unplanted cubes.

HP1b: Glucose-equivalents are higher in planted than in unplanted cubes.

HP1c: Microbial biomass (carbon, bacteria & fungi) is higher in planted than in unplanted cubes.

HP1d: Soil-emitted CO_2_ is higher in planted than in unplanted cubes.

The second set of hypotheses was based on the premises that plants take up nitrogen from the soil and that adding mineral nitrogen fertilizer would compensate plant N-uptake. We expected that the effect of fertilizer addition and plant presence would be visible in soil NO□ concentrations and hypothesized:

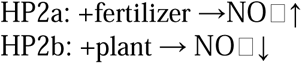

### 2.2 Experimental set-up

The experiment was conducted in the TERRA-Ecotron, where winter wheat (*Triticum aestivium* var. Asory) was grown in the meteorological conditions of the years 2013, 2068, and 2085, representing a gradient of present and predicted climate change scenarios for Central Europe^[10]^. As a second factor, the experiment used two soil types excavated from real farmlands representing historically low (S1) vs. high (S2) soil organic matter inputs (10 vs 22 g SOC kg^-1^ soil respectively, both C:N=10.5). The TERRA-Ecotron has six controlled environment rooms (CERs), in each of which nine cubes (125L) with intact soil monolith can be placed^[14]^. In each CER, eight soil monoliths were planted and one was left unplanted.

Given that two soil types were used, there was one unplanted cube per soil type per climate. In each CER, one cube with plants and the one existing cube without plants were selected for the here presented decomposition study, covering all six soil x climate modalities in planted and unplanted state. For the main study factor of this experiment, the presence or absence of plants, there were thence n=6 replicates of each modality (plant yes/no), comprising the two soils and three climates, which were not replicated within the plant treatment. In each of these cubes, five bags of each tea type (green tea and rooibos tea) were placed at a depth of 8cm (supplementary material SM0), leading to a total of n=120 tea bags being deployed. The study started at the end of winter in 2068 and lasted from DAS131-DAS159. During this time, the wheat was at the beginning, middle and end of tillering in 2013, 2068, and 2085 respectively (approx. BBCH 16,27,38). Only 2085 had received mineral fertilizer at the time of study (Figure 1).

**Figure 1:**
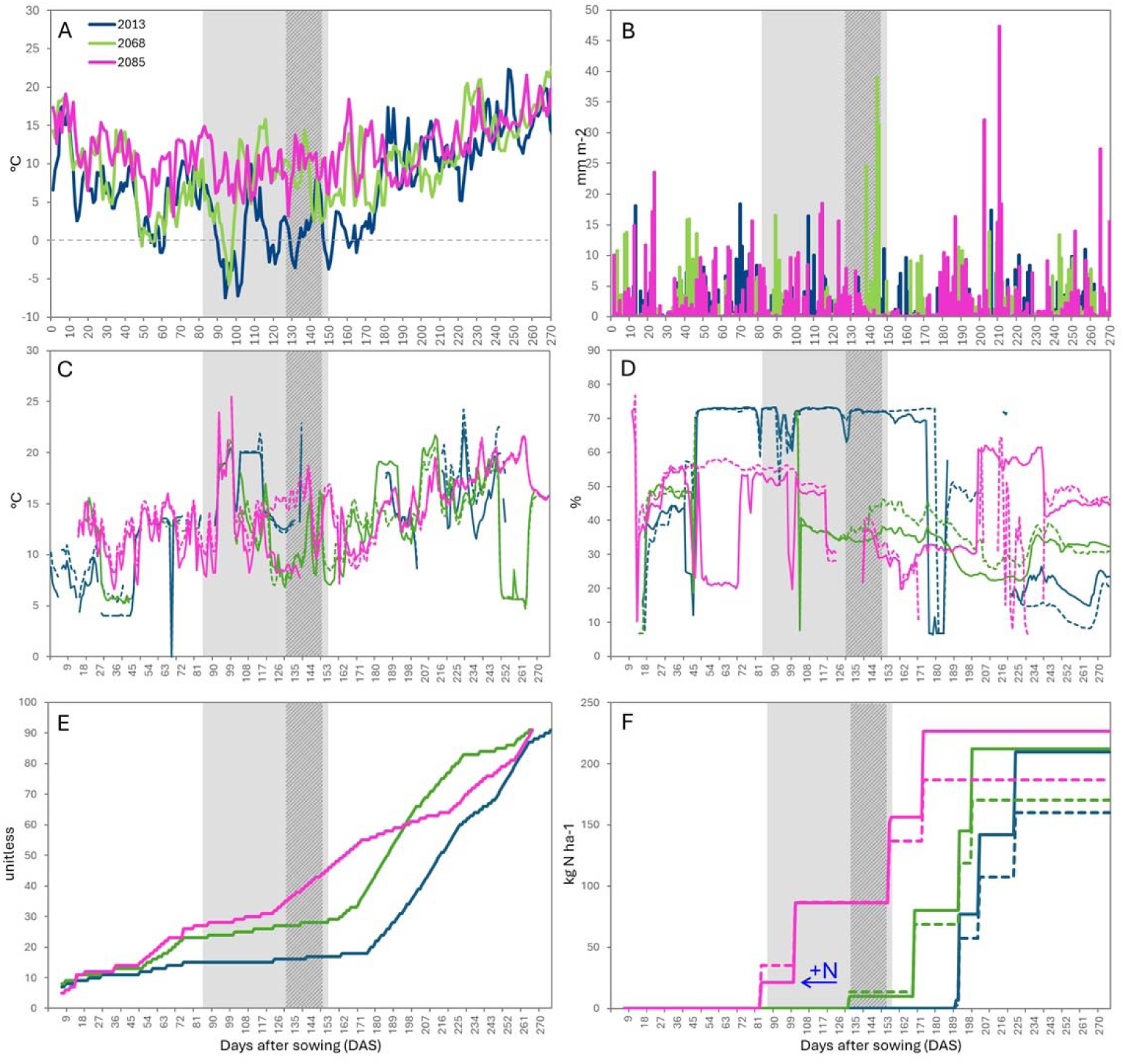
Environmental and soil conditions during the experiment. The study period is shade d in grey in each plot, with the tea bag deployment period (DAS131-145) in dark dashed grey. If individual measures were performed for the two soil types S1 (low OM) and S2 (high OM), S1 is plotted with a straight line and S2 is plotted with a dashed line. A: Daily mean air temperature (°C), B: Daily cumulative precipitation (mm m^-2^), C: Daily soil temperature (°C), D: Daily mean volumetric soil humidity (%), E: Plant growth stages (BBCH), F: Cumulative nitrogen input from mineral fertilizer (kg N ha^-1^). Note: N was applied in three doses according to phenology. Only 2085 had received N at the time of study (at DAS 102), with respective N-application indicated as “+N” in panel F.

### 2.3 Soil and climate conditions during the experiment

The experiment used two soil types of identical soil classification (silt loam (Aba(b)0)) but S2 had received twice as much organic matter than S1, resulting in 2x higher carbon and nutrient contents in S2 at similar C:N and pH (see [10] for detailed soil description). During the tea bag incubation (dark grey area in Figure 1), the air temperature followed the chronological sequence 2013<2068<2085, which also applied to soil temperature where it was measurable 2068<2085, with S2 being warmer than S1 (Table 1). Precipitations followed an inverse chronological pattern of 2013>2068>2085. However, volumetric soil moisture content was similar in 2068 and 2085 reflecting the prior rain input (Figure 1B). Similar to soil temperature, volumetric soil moisture content was also higher in S2 than in S1 in both climates. In S1, the relative soil humidity as determined by drying & weighting of soil subsamples at DAS131 and DAS159 was lower in planted soil than in unplanted soils. This was also the case for S2 in 2085, but in 2068 and 2013 planted S2 soil was wetter than unplanted S2 soil. The pattern was consistent at both DAS131 and DAS159 (SM1).

**Table 1:** Soil temperature and water content during the time of tea bag incubation (DAS 131-145). ‘Y’ and ‘NO’ indicate planted (YES) and unplanted (NO) soil respectively. See SM1 for soil moisture data separated for DAS131 and DAS159. *WR: winter room: no data from soil sensors.

|  | 2013 |  |  |  | 2068 |  |  |  | 2085 |  |  |  |
| --- | --- | --- | --- | --- | --- | --- | --- | --- | --- | --- | --- | --- |
|  | S1 |  | S2 |  | S1 |  | S2 |  | S1 |  | S2 |  |
| Air T □ C | 2.15 |  |  |  | 7.91 |  |  |  | 10.72 |  |  |  |
| Soil T □ C | WR* |  | WR* |  | 10.15 |  | 10.56 |  | 11.77 |  | 16.20 |  |
| Δ soil T □ C S2-S1 | NA |  |  |  | +0.41 |  |  |  | +4.43 |  |  |  |
| Cumulative precipitation mm m<sup>-2</sup> | 180.40 |  |  |  | 136.24 |  |  |  | 13.25 |  |  |  |
| Vol. soil moisture content (MC) % | WR* |  | WR* |  | 34.19 |  | 38.47 |  | 34.45 |  | 37.99 |  |
| Δ soil vMC % S2-S1 | NA |  |  |  | +4.27 |  |  |  | +3.54 |  |  |  |
| Rel. soil MC % | Y | NO | Y | NO | Y | NO | Y | NO | Y | NO | Y | NO |
| Av. DAS131-159 | 18.91 | 21.65 | 21.92 | 10.34 | 17.84 | 20.72 | 19.77 | 17.85 | 12.81 | 16.74 | 15.35 | 19.64 |
| Δ soil vMC % unplanted-planted | +2.74 |  | -11.58 |  | +2.88 |  | -1.92 |  | +3.93 |  | +4.29 |  |

### 2.4 Decomposition and further carbon and nitrogen cycling parameters

Within each of the studied cubes, n=5 rooibos and n=5 green tea bags were buried at 8cm depth to measure decomposition with the tea bag index from DAS131-145. The tea bag index integrates mass loss data from both green tea (fast-decomposing) and rooibos tea (slow-decomposing) into a single standardized k value combining information from the two substrates to represent the early-stage decomposition rate under asymptotic model assumptions (SM8)^[15]^. The tea bag index is most indicative for litter decomposition, but tea is mineralized alongside soil organic matter and plant-derived compounds if present. In addition, in each cube interstitial soil pore water was extracted in situ using permanently installed soil pore water samplers (Rhizosphere Research Products B.V., Wageningen, The Netherlands) with the 10cm long porous membrane situated in the center of each cube vertically inserted at a depth of 8-15cm. Water was extracted weekly (DAS 82, 102, 110, 131, 150) by applying a negative pressure using 20 ml syringes and metabolically relevant compounds were quantified immediately after water extraction using anthrone reaction for glucose equivalents (modified after [16]) and nitrate was quantified biochemically using an ion-selective electrode (LAQUAtwin NO3-11C, Horiba Ltd., Kyoto, Japan). At DAS 102, 131 and 159, basal as well as exudate-induced respiration were quantified via MicroResp with two technical replicates for each modality^[17]^. Soils were sieved to 2 mm and incubated within the CERs without further adjusting water content to maintain the soil samples as close to the humidity and temperature conditions of respective climate scenarios. Basal respiration was measured as CO_2_ released from soil samples without any amendment, and as a proxy for exudate-induced respiration, 25µl of the collected interstitial soil pore water was used. At DAS131 and DAS159, microbial biomass and bacterial and fungal gene copies were measured in n=4 planted cubes per CER and the n=1 unplanted cube using direct chloroform extraction followed by titration (modified after [18]) and qPCR of 16S and 18S respectively (SM 2)^[19,20,21]^. Sensors for continuous monitoring of soil humidity and temperature were installed in n=2 planted cubes per soil type per CER, but was not permanently monitored in unplanted cubes. Therefore, soil humidity was also quantified using the gravimetric method, drying sub-samples to constant mass, for all soils of planted and unplanted cubes which were also used for microbial biomass and qPCR at DAS131 and DAS159.

### 2.5 Statistical analysis

The individual and interactive effects of plant (YES,NO), climate (2013,2068,2085), soil type (S1,S2), and DAS (82,102,110,131,150) where applicable, on the response parameter decomposition, glucose, nitrate, microbial biomass, fungal and bacterial genes, and basal and exudate-induced respiration were individually evaluated with linear mixed models, with a random effect for cube (y∼ PLANT * (SOIL + YEAR + DAS) + (1 | CUBE)).

To increase the resolution of the potential driving factors of decomposition (k) beyond the fixed effects climate (temperature, rain), soil (SOC, sand, humidity, temperature), plant (total root length, network area, branching frequency, number of root tips) and rhizosphere processes (glucose, nitrate, microbes), partial least squares regression (PLS) was performed^[22]^. Leave-one-out cross-validation was used to assess model performance and the orthogonal scores algorithm was selected, as required for subsequent calculation of variable importance in projection (VIP) scores (SM3, SM4). Leave-one-out cross-validation (LOO-CV) was employed to assess model performance and determine the optimal number of components. Missing values in the predictor matrix were imputed using median values per variable, and constant predictors were removed prior to scaling and centering the data. To complement the PLS analysis and capture potential non-linear relationships and interactions among predictors, a random forest (RF) regression model was fitted^[23]^. The model used 500 trees and the default number of variables tried at each split for regression (mtry = floor(p/3), where p is the number of predictors). The same imputed predictor matrix (without scaling) and response variable (k) were used as in the PLS model. Model performance was evaluated via out-of-bag (OOB) mean squared error and % variance explained (SM5, SM6). Both methods were applied to the same pre-processed dataset to enable direct comparison of predictor importance rankings. Overlap in the top-ranked variables across PLS (VIP > 1) and random forest (%IncMSE) provides robust support for the key drivers of decomposition.

Data was analyzed using statistical software R^[24]^, with the additional packages car^[25]^, caret^[26]^, dplyr^[27]^, effects^[28]^, emmeans^[29]^, ggnewscale^[30]^, ggplot2^[31]^, lme4^[32]^, lmertest^[33]^, patchwork^[34]^, performance^[35]^, pls^[36]^, plsVarSel^[37]^, randomForest^[38]^, tibble^[39]^, tidyr^[40]^, and tidyverse^[41]^.

## 1. Results

### 3.1 Decomposition constant k

Plant presence significantly influenced the decomposition constant k (F□,□ = 10.64, p = 0.031), with higher values in unplanted (NO) compared to planted (YES) cubes when averaged across soil types and climate scenarios (Figure 2). A marginal main effect of climate scenario was also observed (F□,□ = 6.27, p = 0.058), whereas soil type had no significant effect (F□,□ = 0.21, p = 0.674). No significant interactions were detected between plant presence and soil type (PLANT × SOIL: F□,□ = 3.76, p = 0.125) or between plant presence and year (PLANT × YEAR: F□,□ = 4.48, p = 0.095), indicating that the negative effect of plant presence on decomposition rate was relatively consistent across soil types and climate conditions. However, there was a tendency for higher decomposition in S2 than in S1 in unplanted soils, most notably in 2013, and higher decomposition in S1 than in S2 in presence of plant, most notably in 2085.

**Figure 2:**
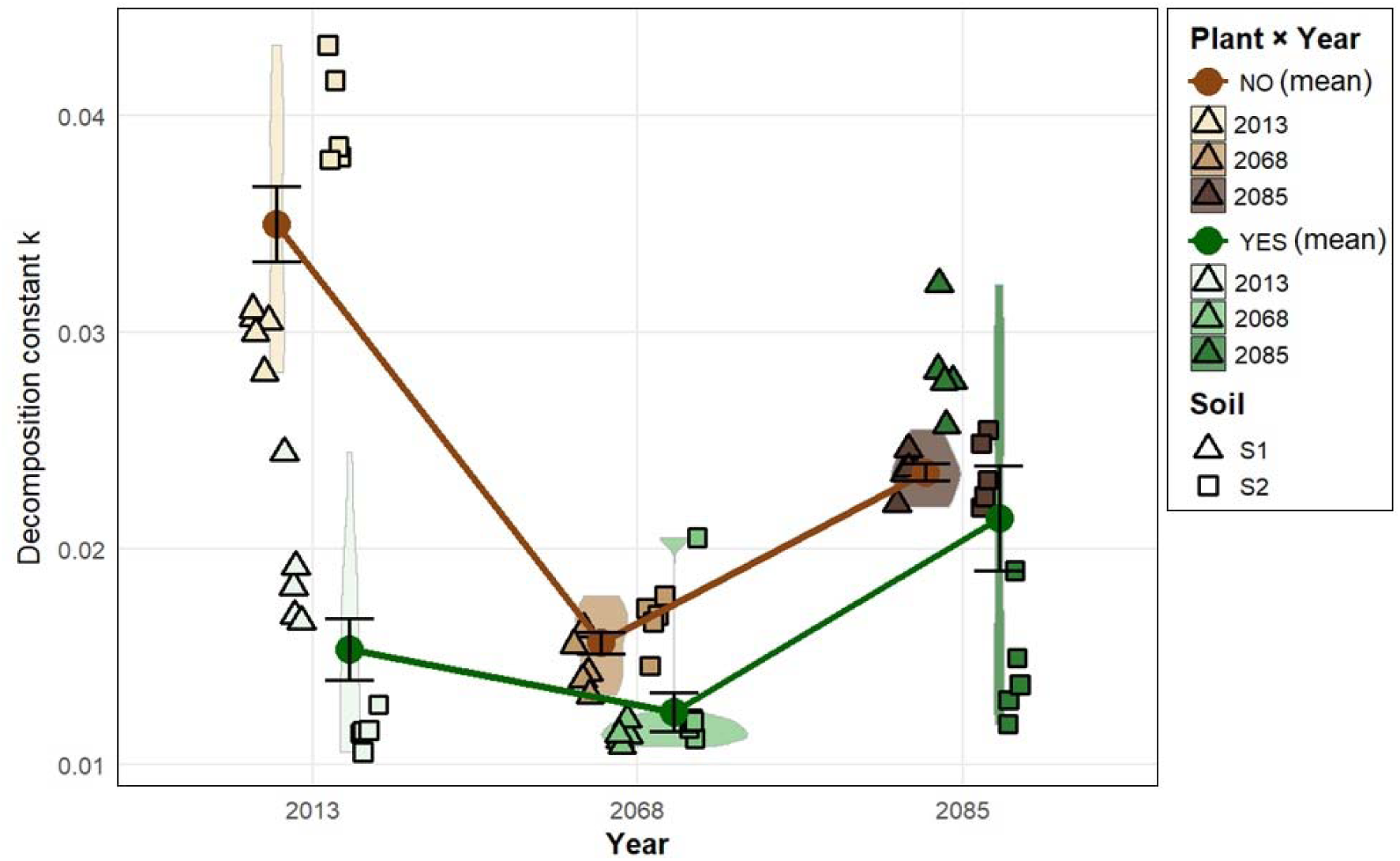
Decomposition constant k as determined by the tea bag index in planted (YES) and unplanted (NO) soils in three climates (2013, 2068, 2085) under two soil management histories (S1: low OM, S2: high OM). Triangles and squares represent the technical replicates (individual tea bags) within each soil x climate modality (n=5) and circles and lines represent respective means.

### 3.2 Glucose-equivalents and nitrate in interstitial soil pore water

For glucose levels in interstitial soil pore water, linear mixed-effects models revealed a significant interaction between plant presence and days after sowing (PLANT × DAS: F□,□□□ = 13.69, p <0.001), where planted cubes exhibited lower soil glucose concentrations than unplanted controls at most time points (Figure3 upper panel). No significant interactions were found between plant presence and soil type (PLANT × SOIL: F□,□ = 1.46, p = 0.293) or climate scenario (PLANT × YEAR: F□,□ = 0.049, p = 0.953), suggesting that the plant-induced reduction in glucose was consistent across soil types and climate conditions. There was an overall decline of glucose concentration in interstitial soil pore water by -0.0045 units per 10 days from DAS82 to DAS150, with a spike at DAS131. When averaged across all sampling dates, the main effect of plant presence was not significant (p = 0.110), and neither soil type (p = 0.720) nor year (p = 0.439) had significant main effects.

Plant presence also had a significant overall effect on nitrate levels which were lower in planted than in unplanted cubes (F□,□ = 9.70, p = 0.036), most likely indicating plant N-uptake (Figure 3 lower panel). This effect was modulated by sampling time, as indicated by a significant PLANT × DAS interaction (F□,□□□ = 98.14, p < 0.001). A marginal interaction between plant presence and climate scenario was also observed (PLANT × YEAR: F□,□ = 6.16, p = 0.060), whereas no significant interaction with soil type was detected (PLANT × SOIL: F□,□ = 0.063, p = 0.814). Neither soil type (p = 0.496) nor year (p = 0.565) exerted significant main effects.

**Figure 3:**
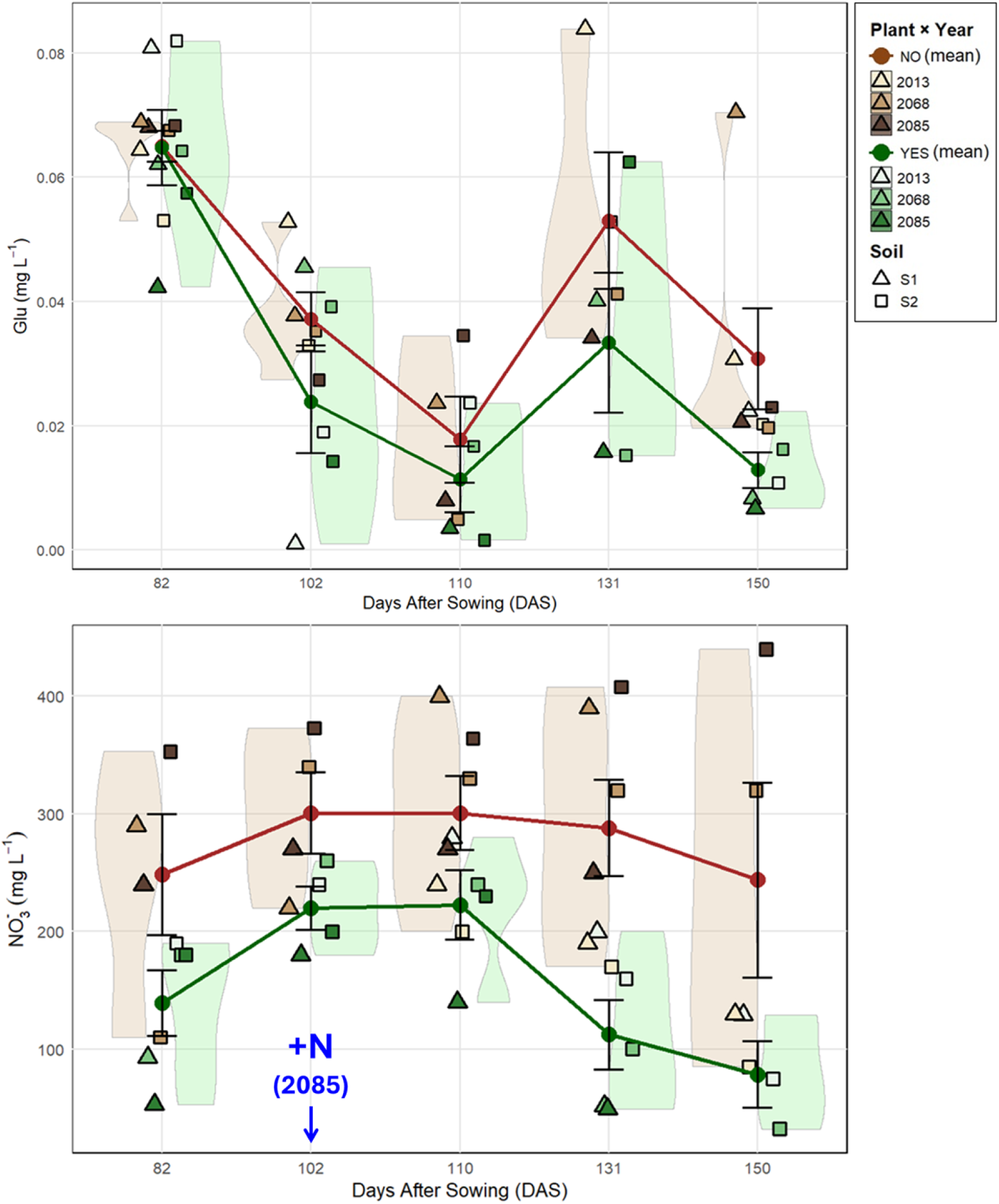
Glucose (upper panel) and nitrate (lower panel) dynamics in interstitial soil pore water determined at five days (days after sowing (DAS)). Triangles and squares represent one value per soil x climate modality, totaling between four and six values for each level of the main factor ‘plant’ (planted=YES and unplanted=NO) at each day, depending on the extractability of interstitial soil pore water. Note: Mineral nitrogen fertilizer (NO_3_NH_4_) was applied at DAS102, only in 2085 and only in planted cubes.

### 3.4 Microbial biomass carbon, bacteria and fungi

No significant main effect of plant presence on bacterial gene copy numbers, fungal gene copy numbers or microbial biomass carbon (mbC) was detected at 131DAS and 159DAS (all p > 0.43). Time (DAS) also had no significant effect on bacterial gene copies (F□,□□ = 0.071, p = 0.796), fungal gene copies (F□,□ = 2.23, p = 0.170) and microbial biomass C (F□,□□ = 0.239, p = 0.635). Similarly, no significant interactions were detected in any variable and neither soil type nor year showed statistically significant effects. However, on average the measured parameter were higher in unplanted than in planted soils, apart from mbC at 159DAS.

**Figure 4:**
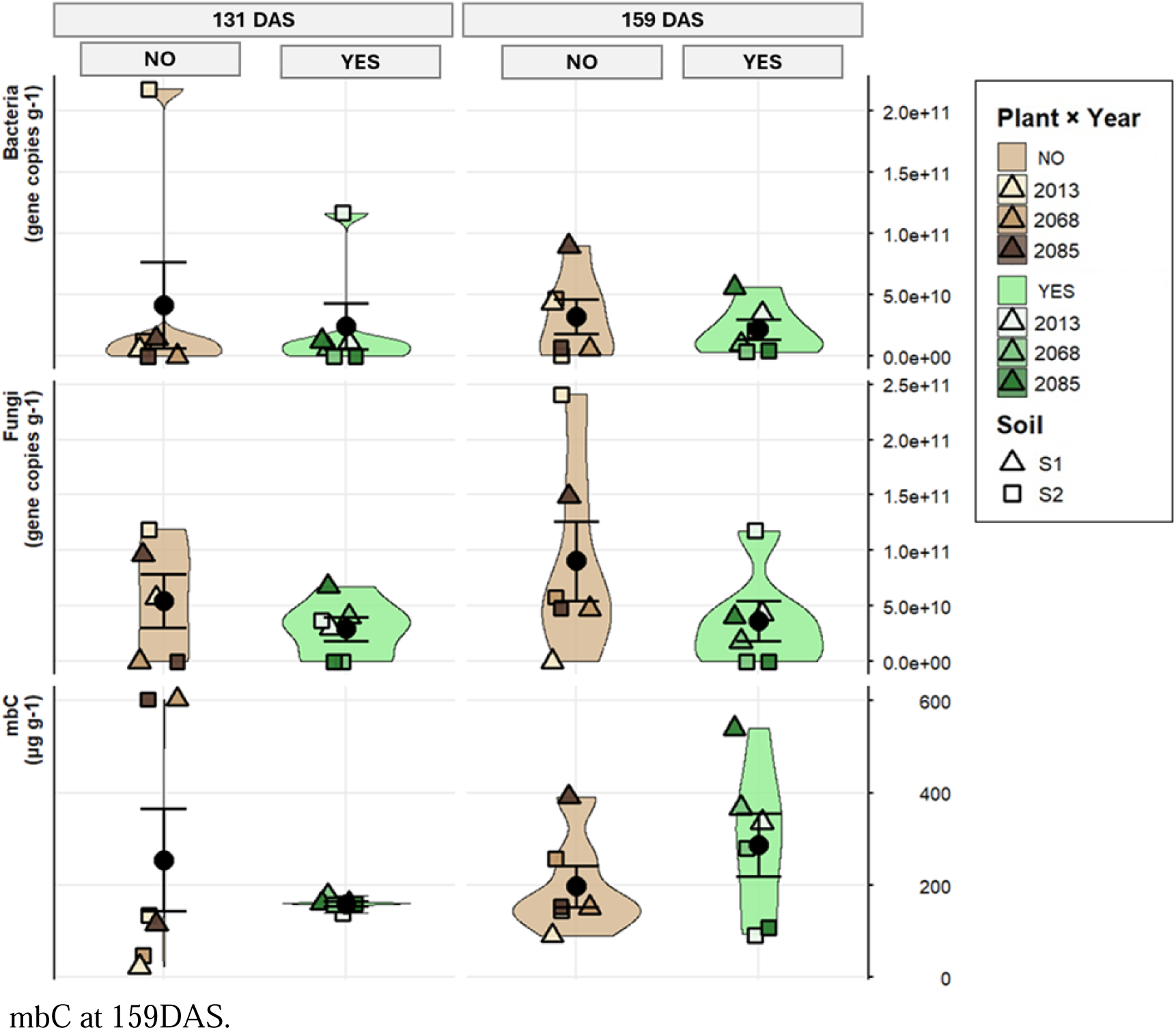
Bacteria (16S gene copies g soil ^-1^) and fungi (18S gene copies g soil ^-1^) abundance and microbial biomass carbon (mbC g soil ^-1^) at two time points (DAS131, DAS159) in unplanted (NO) planted (YES) cubes in two soil types (S1: low OM, S2: high OM) in three climates (2013, 2068, 2085). Triangles and squares represent one value per soil x climate modality, totaling six values for each level of the main factor ‘plant’ (planted=YES and unplanted=NO) at each day.

### 3.5 Basal and exudate-induced soil respiration

Repeated-measures linear mixed models with random intercept for “cube” were fitted separately to basal and exudate-induced CO□ data. There was no overall significant effect of plant on either CO_2_ parameter, but both CO_2_ parameters were higher in the future climates than in the past climate.

Specifically, basal CO□ showed no significant influence of plant presence (main effect: F□,□□ = 0.168, p = 0.686) and no significant interaction with time (PLANT × DAS: F□,□□ = 0.133, p = 0.877). Significant effects were observed for DAS (F□,□□ = 10.95, p = 0.00055) and year (F□,□□ = 4.80, p = 0.019) but not soil type (p = 0.802).

Exudate-induced CO□ likewise showed no significant main effect of plant (F□,□□ = 0.317, p = 0.580) or PLANT × DAS interaction (F□,□□ = 0.216, p = 0.807). Time (DAS: F□,□□ = 6.80, p = 0.0053) and year (F□,□□ = 4.43, p = 0.025) were significant, whereas soil type had no effect (p = 0.245).

### 3.6 Integrative analysis

Despite several climate, soil and root parameter being included, the most powerful predictors of decomposition were microbial and metabolic factors, above all fungal gene copies and microbial respiration (CO_2_) which was constituent for both RF and PLS models (Figure 6).

**Figure 5:**
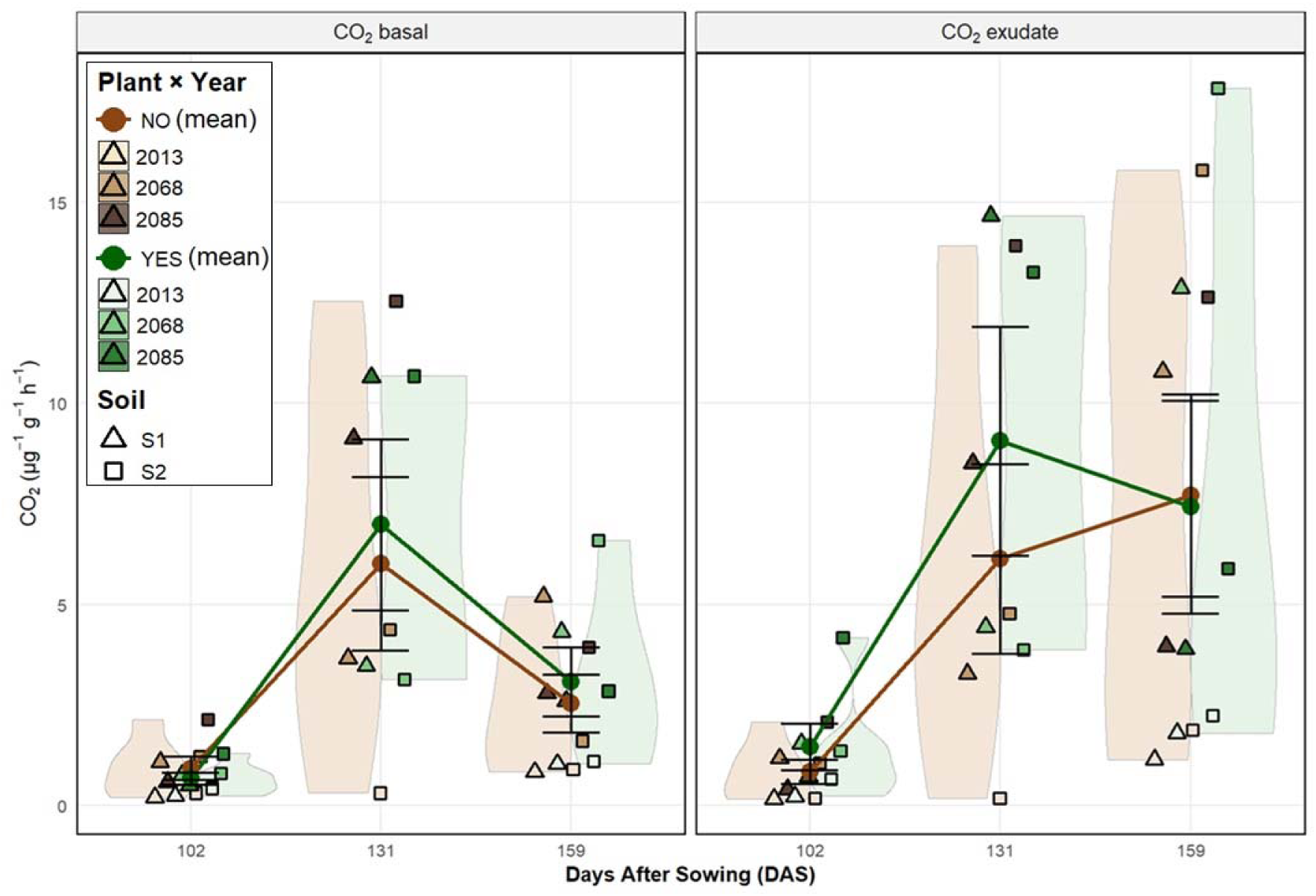
Basal (left) and exudate-induced (right) soil respiration (µg CO_2_ g soil ^-1^ hour ^-1^) in planted and unplanted cubes in two soil types (S1: low OM, S2: high OM) in three climates (2013, 2068, 2085) at three timepoints (DAS102, DAS131, DAS159). Circles and triangles represent one value per modality, each of which is the mean of two technical replicates. This totals six values for each level of the main factor ‘plant’ (planted=YES and unplanted=NO) at each day.

**Figure 6:**
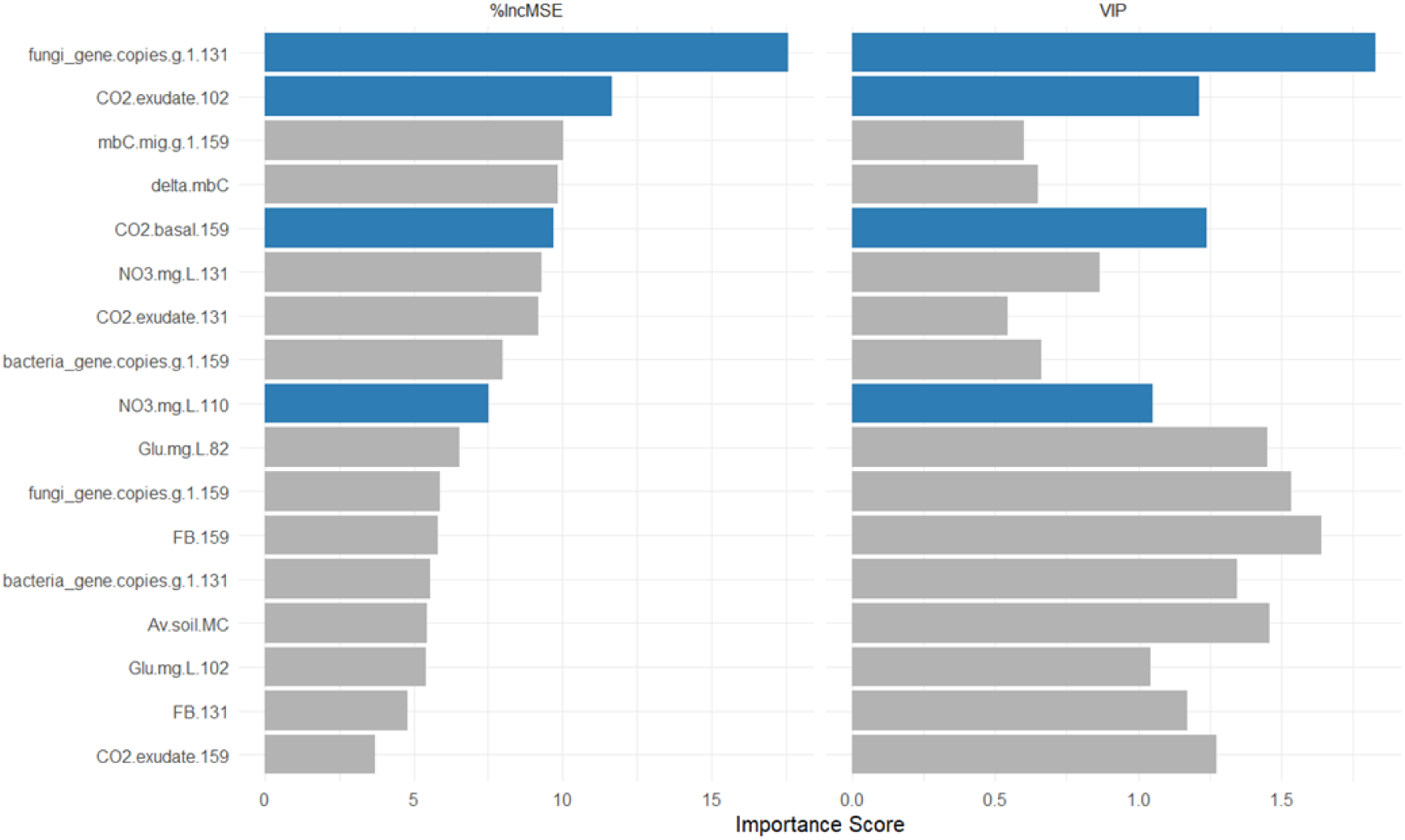
Variable importance comparison for predicting the decomposition constant k using Random Forest (RF) regression (left) and Partial Least Squares (PLS) regression (right). Bars represent importance scores for predictors meeting at least one of the following criteria: Variable importance in projection (VIP in PLS) > 1 or percentage increase in out-of-bag mean squared error when the variable is randomly permuted (%IncMSE in RF) > 7 (breakpoint cf SM6). Variables which meet both criteria are highlighted in blue.

The most significant environmental variable was average soil moisture content (Av.soil.MC). Strong agreement between the two independent methods supports notably the importance of fungal gene copies at 131 DAS, alongside consistent high rankings for nitrate (NO□) concentrations, CO□ respiration (basal and exudate-induced), microbial biomass carbon (mbC), and related rhizosphere indicators at later sampling points. Partial dependence plots (SM7) revealed strong positive associations between the decomposition constant k and fungal gene copies at 131DAS, nitrate concentrations at 131DAS, microbial biomass carbon at 159DAS and glucose at 150DAS, indicating these as key drivers of decomposition. In contrast, negative associations were observed for exudate-induced CO□ respiration at 102DAS and basal CO□ respiration at 159DAS (SM7). Nonetheless, the proportion of variance explained by RF was substantially higher than by PLS (95% vs 18%) indicating that interactions among the included variables are also important.

## 4. Discussion

### 4.1 Plant-microbe competition (hypotheses under premise 1)

Premise P1 posited that plant exudates of carbohydrate-rich substances enhance microbial mineralization^[1,2,3,4]^. Contrary to this, results indicate that plant presence slowed rather than amplified decomposition (Figure 2), which has been observed previously for grasses and forbs^[42]^. The lower decomposition in planted than in unplanted cubes was in line with lower glucose and nitrate concentrations in planted cubes, leading to rejection of HP1a and HP1b. Under the given measurement conditions and time frame, no overall differences emerged in soil-emitted CO□, microbial biomass, or bacterial/fungal gene copies, leading to rejection of HP1c and HP1d. These results align with resource competition theory between plants and microbes, which posits that plants can outcompete microbes over nutrient acquisition, inducing N-limitation for microbes and thus delaying decomposition^[5,7,43,44,45]^. Every molecule of nitrogen which goes into belowground biomass of microbes or fauna, is a molecule of nitrogen which is not immediately available for plant uptake. Microbes thus often dominate short-term N-competition, but over longer timeframes competition for nitrogen is often asymmetrical in favor of the plants, sometimes accompanied by positive rhizosphere priming effects that enhance microbial SOM-mineralization^[1,46,47,48]^. This asymmetry is context-dependent, but ¹□N-labeling studies indicate that plants ultimately acquire more fertilizer-derived N than microbes, particularly in the rhizosphere^[6,49]^.

Implementing such dynamics into concepts of "soil health" is important, as high microbial abundance and biomass can present a significant N-sink and thus antagonize plant growth ^[5,10,50]^. Whether or not belowground organisms benefit plant growth depends on food-web structure, on the type of relationship established between symbiotic organisms and plants (mutualism, commensalism, parasitism), and whether catabolism or anabolism prevails in soil organisms^[51,52]^. To optimize interactions between crops and soil organisms for improved plant production, future research could identify parameters which allow to distinguish facilitative and antagonistic organisms, including links to carbon and nutrient exchange dynamics^[4,53,54,55,56]^.

### 4.2 Nitrate homeostasis (hypotheses under premise 2)

Under premise P2 we assumed that plants take up nitrogen and that adding mineral nitrogen fertilizer compensates or even exceeds plant N-uptake and thence expected to observe an increase in nitrate concentration in interstitial soil pore water in 2085 (or a less steep decrease). The addition of fertilizer at DAS102 in 2085 did not result in measurable differences in interstitial soil pore water, which increased also in the unfertilized planted soils and in the unplanted soils at that moment of measurement, leading to rejection of H2a (Figure 3). While we observed lower nitrate concentrations in planted than in unplanted soils, we reject H2b at α=0.05, despite robust indication for substantial plant N-uptake (average depletion −91 mg L^-1^, p = 0.08).

Similarly, there was no indication that warmer temperatures in future climate scenarios stimulated microbial activity during the study period, accelerating decomposition and thus enhancing interstitial soil pore water. While this observation is in line with other studies^[57,58]^, the absence of a detectable effect of fertilizer addition or temperature on soil NO□ does not constitute evidence of absence of such effects, as this result might have been caused by the measurement conditions, where rapid plant and microbial N-uptake could have obscured the performed measurements in interstitial soil-pore water. Plants exhibit high-affinity uptake systems for NO□□, allowing quick absorption from soil solution especially during periods of active growth like the tillering stage in cereals^[59]^. Over time, soil nitrogen concentrations could also be stabilized as a function of the net balance of nutrient inputs and outtakes in the belowground multi-trophic networks, also known as the ’microbial loop,’ wherein the predation of bacteria and fungi by microbial grazers, such as protozoa and nematodes, triggers the mineral release of previously immobilized nitrogen back into the rhizosphere^[60,61]^.

In addition, variable precipitation, especially reductions or erratic patterns, can compound nutrient losses by drought-stress induced disruptions of microbial community structures and reduces enzyme activities involved in nutrient retention, potentially leading to higher losses of both carbon and nitrogen^[58,62,63]^. In such scenarios, soils with diverse but less resilient microbial communities, for example those dominated by fast-growing opportunists, may experience amplified nutrient volatilization or erosion, particularly in agricultural or grassland systems^[64,65]^.

### 4.3 Role of stoichiometry

Reduced decomposition has been previously observed for various grasses and forbs and it has been proposed that this is driven by elevated C:N ratios of root exudates, providing microbes with abundant energy but insufficient nitrogen^[42,66]^. This stoichiometric imbalance can trigger microbial nitrogen sequestration (immobilization), when microbes prioritize the uptake of scarce inorganic N like nitrate and ammonium to support their own biomass growth^[6]^. When the substrate to be decomposed is also nutrient-poor with high C:N ratios, which is the case for organic matter from plant residues like straw and tea bag litter, it can result in enhanced competition for nitrogen between plants and microbes leading to a net decrease in decomposition rates due to severe nutrient limitation within the microbial community. The typical C:N ratio of wheat straw ranges from 80:1 to 100:1, though it can occasionally be as high as 120:1 depending on the variety and fertilization history^[67]^.

Because these values vastly exceed the critical C:N threshold of approximately 25:1 to 30:1, microbial demand for nitrogen surpasses the supply available in the residue, leading to net N sequestration (immobilization) in the microbial biomass rather than mineral release^[68]^. Green Tea has a C:N of approximately 10:1 to 12:1, which is below the critical threshold (< 25:1). Thus, it undergoes faster mineralization than rooibos tea, which has a C:N of approximately 40:1 to 60:1 and is thus more prone to trigger nitrogen competition (SM8)^[15,69]^.

### 4.4 Fungi and humidity are more important for decomposition than bacteria and temperature

In multivariate analysis including air and soil temperature and precipitation and soil moisture, as well as several soil and root parameters, microbial and metabolite parameters remained the strongest in predicting decomposition, in particular fungal gene copies and CO_2_ as an indicator of microbial activity. Fungi are generally associated with the breakdown of more complex molecules, while bacteria are thought to feed primarily on LMW sugars, which is in line with this result^[70,71]^. However, these roles are highly context-dependent. For example, some rhizosphere fungi can outcompete bacteria for labile exudates under nutrient-limiting conditions, and selective grazing from protists and nematodes can also shift fungal-to-bacterial ratios thereby altering decomposition and N-availability^[72,73]^. However, there was no interaction effect with nitrogen addition in 2085, which has previously been shown to affect wheat straw decomposition through altered bacteria and fungi dynamics^[74]^. Soil moisture was the only environmental parameter having a considerable influence on the model’s predictive power (Figure 6). Soil moisture influences decomposition and microbial activity by regulating oxygen availability and substrate diffusion^[75,76]^, which possibly contributed to the increased decomposition rates particularly in unplanted cubes in 2013, which was the wettest year during the study period (Figure 1). Decomposition is generally considered to be also temperature sensitive^[77]^. Temperature’s lack of relevance in this multivariate context could have stemmed from its effects being constrained or masked by other factors such as moisture or microbe-substrate interactions^[78]^. On the other hand, previous studies have found that microbial decomposition can be very resistant to temperature^[79,80]^, so the absence of temperature effects on decomposition could also indicate true absence of ecological relevance in this context.

### 4.5 Possible impact of winter conditions on soil processes in 2013

The biggest difference in decomposition between planted and unplanted cubes was observed in the 2013 climate, which was also the year where plants and soils experienced winter conditions with below zero temperature (Figure 1, Figure 2). Freeze-thaw cycles can disrupt microbial cells and soil aggregates, leading to cell lysis and the release of organic matter, including labile carbon and nitrogen compounds^[81]^. This "thaw burst" can enhance microbial mineralization rates as surviving or recolonizing microbes rapidly decompose the newly available substrates, increasing nutrient cycling and CO□ emissions in the short term^[82]^. In the unplanted soils, all elements released through this process were available to microbes, increasing decomposition. In the planted soils, plants also took up some of the nutrients released by microbial turnover, thus lowering resources for microbes and therewith their metabolic activity in mineralizing organic materials. Prolonged freezing can also reduce overall microbial biomass and diversity over winter, delaying mineralization recovery^[83]^. If thaw coincides with erratic precipitation, it can lead to nutrient losses via leaching and/or denitrification, reducing net mineralization efficiency, which aligns with the lower overall NO_3_^-^ and respiration in 2013 (Figure 3, Figure 5). Microbes often outcompete plants for N post-thaw due to higher surface area and rapid growth, leading to temporary N-immobilization and suppressed plant uptake^[84]^. This could explain the lower decomposition, and NO□□ in planted soils in 2013 as plants can exacerbate resource scarcity. In unplanted soils, microbial activity is not limited by resource drain from plant nutrient uptake, possibly amplifying differences between planted and unplanted soils. Warmer winters in the future could reduce the nutrient flush from thawing events, leading to more balanced nutrient pools but potentially higher overall microbial N-sequestration if their activity is not limited by carbon or other factors.

### 4.6 (No) differences between soil types

In the low organic matter soil S1, the unplanted soil was wetter than the planted, most likely due to plant water-uptake in the planted cubes^[85]^. Interestingly, in the high organic matter soil S2, the unplanted soil was drier than the planted soil in 2013 and 2068, possibly due to increased drainage and/or evapotranspiration in unplanted S2 cubes, and/or due to increased water-holding capacity of planted cubes in S2 in these years^[86,87]^. Decomposition rates were higher in unplanted S2 in 2013 and in planted S1 soils in 2085 (Figure 2). This aligns with resource competition dynamics, where plant presence may suppress decomposition in high-SOM soils via nutrient limitations on microbes, while enhancing it in low-SOM contexts through priming effects, though this was not reflected in soil CO_2_-emissions as measured here^[42,46]^. Notably, this mirrors plant performance and yield trends, which were higher in S1 than S2, likely because lower SOM in S1 reduces microbial competition for nutrients, allowing greater crop access and productivity, while higher SOM in S2 supports faster microbial decomposition but may limit yields through antagonistic interactions or nutrient immobilization, in contrast to expectations^[12,10,50,88]^.

Despite these contrasting soil water dynamics between S1 and S2, measured parameters like decomposition, respiration, and glucose equivalents showed minimal differences overall, indicating that this process is either driven by stoichiometry as both soils have a C:N ratio of 10.5, and/or that plants and soil microbes can buffer against water and temperature variability to sustain homeostasis in nutrient cycling and microbial activity^[89,90]^. This resilience aligns with findings of no significant grain quality differences between these two soil types in the same experiment^[91]^, though climate exerted a broad negative impact on grain quality, highlighting the sensitivity of crops to abiotic stressors despite biotic buffering.

### 4.7 Implications for agroecosystems under climate change

The slowing down of decomposition in the presence of growing plants as demonstrated here and elsewhere^[42]^ supports the integration of cover crops in rotations as they can prevent N-leaching and provide habitat for diverse species^[92]^. It has also been suggested that inhibition of decomposition by ecto-mycorrhizal fungi in forested ecosystems resulting from plant-microbe-competition can increase soil C storage^[93,94]^, which could also be a mechanistic explanation for increased soil C-stocks described under cover crops^[95,96]^. However, the results of this study also indicate that soil management and fertilization practices can still be improved to synchronize soil nutrient provision with plant nutrient demand, particularly at critical growth stages like stem elongation. One possible lever to improve N-availability for plants which might become even more relevant under future climates is nitrification inhibition, either synthetic or biological^[97,98]^. Another path towards mitigating the risk of nutrient losses to plant-microbe competition could be increasing the inherent capacity of cereals to acquire nitrogen-fixing microbes^[99]^. Similarly, practices like intercropping or integrating legumes into crop rotations could help balance plant-microbe dynamics and promote steady mineralization while mitigating nutrient losses to competition, however this does not always benefit yields^[100,101]^. The results of this study also reinforce the benefits of split fertilization in cereal systems, as nutrient demands at critical growth stages can be high necessitating readily plant-available nitrogen to support plant growth and fitness, and ultimately yields and protein contents^[102]^.

### 4.8 Limitations & outlook

The present study covered a period of 14 days for the assessment of decomposition via the tea bag index, which is an appropriate timeframe to capture the majority of overall mass loss from tea bags and differences between green and rooibos tea (SM8)^[15]^. Yet we acknowledge that this period represents solely the conditions around the stem elongation growth stage of the wheat plants in the respective climates. Decomposition dynamics may be different at different growth stages^[47,103]^ which were not studied in the present experiment to avoid confounding factors such as diverting growth stages due to staggered development and differences in external inputs from multiple doses of fertilization (Table 1, Figure 2). Further, given that microbial processes and plant N-uptake can be fast (seconds-hours), it is possible that some of the dynamics were missed despite the methods used being very sensitive.

Finally, the main factor of this experiment was plant presence or absence, for each of which we had six independent replicates, but nested within the main factor ‘plant’ were the factors soil and climate for which only technical replicates exist. While we can have confidence in the main plant-effect which was suppression of decomposition consistent across all soil x climate modalities and in line with previous studies^[42]^, future experiments could increase the replication for nested factors like soil type and climate to further nuance the findings. Future studies could also increase the spatial and temporal resolution of data acquisition e.g. by increasing rates of sampling for highly dynamic parameter, installing continuous nitrate sensors and potentially tracking C and N molecules using stable isotopes.

## 5. Conclusion

Overall, this study supports the inclusion of plant-microbe competition and of the sink strength of belowground networks into nutrient cycling models for the rhizosphere. By prioritizing approaches that account for these interactions, crop production can be tailored towards reliable soil nutrient provision and adapted to altered plant nutrient demand under varying meteorological conditions. Future research with broader temporal scales could further define these insights for different plant growth stages. Quantifying the sink strength of belowground trophic networks could also inform the timing of fertilizer use to reduce the risk of nutrient losses especially when plant demand is high. Future management strategies could leverage the observed plant-microbe dynamics to enhance the ability of the microbial loop to stabilize nitrogen against leaching during extreme weather while simultaneously improving the synchrony between mineralization and the spatio-temporal nutrient demands of crop plants. By optimizing this biological buffer system, agroecosystems can achieve greater climate resilience, maintaining both soil health and productivity under increasingly volatile environmental conditions.

## Supporting information

Supplementary meterial for tea time

## Data availability

All data presented in this manuscript is available at https://doi.org/10.6084/m9.figshare.31607071.

## Competing interests

The authors have no competing interests to declare.

Ethics, Consent to Participate, and Consent to Publish declarations: not applicable.

## References

1 Dijkstra, F. A., Carrillo, Y., Pendall, E., & Morgan, J. A. (2013). Rhizosphere priming: a nutrient perspective. Frontiers in microbiology, 4, 216. 10.3389/fmicb.2013.00216.

2 Keiluweit, M., Bougoure, J. J., Nico, P. S., Pett-Ridge, J., Weber, P. K., & Kleber, M. (2015). Mineral protection of soil carbon counteracted by root exudates. Nature Climate Change, 5(6), 588–595. 10.1038/nclimate2580.

3 Liu D, Xu L, Wang H, Xing W, Song B, Wang Q. (2024). Root Exudates Promoted Microbial Diversity in the Sugar Beet Rhizosphere for Organic Nitrogen Mineralization. Agriculture 14(7):1094. 10.3390/agriculture14071094.

4 Shabtai, I.A., Hafner, B.D., Schweizer, S.A., Höschen, C., Possinger, A., Lehmann, J. & Bauerle, T. (2024). Root exudates simultaneously form and disrupt soil organo-mineral associations. Commun Earth Environ 5, 699. 10.1038/s43247-024-01879-6.

5 Hodge A, Robinson D, Fitter A. (2000). Are microorganisms more effective than plants at competing for nitrogen? Trends Plant Sci.5(7):304–8. 10.1016/s1360-1385(00)01656-3.

6 Kuzyakov, Y. and Xu, X. (2013). Competition between roots and microorganisms for nitrogen: mechanisms and ecological relevance. New Phytol, 198: 656–669. 10.1111/nph.12235.

7 Zhu, Q., Riley, W. J., Tang, J., and Koven, C. D. (2016). Multiple soil nutrient competition between plants, microbes, and mineral surfaces: model development, parameterization, and example applications in several tropical forests, Biogeosciences, 13, 341–363, 10.5194/bg-13-341-2016.

8 L’Espérance E, Bouyoucef LS, Dozois JA, Yergeau E. (2024). Tipping the plant-microbe competition for nitrogen in agricultural soils. iScience 27(10):110973. 10.1016/j.isci.2024.110973.

9 Jägermeyr, J., Müller, C., Ruane, A.C. et al. (2021). Climate impacts on global agriculture emerge earlier in new generation of climate and crop models. Nat Food 2, 873–885 10.1038/s43016-021-00400-y.

10. Michel J, Leemans V, Weinmann M, Bin J, Biver S, Blum A, Börger R, Cao D, Him SL, Kirbas G, Le Gouis J, Moya-Laraño J, Persyn M, Quenon A, Sanchez-Moreno S, Van Der Straeten D, Waibel M, Xayphrarath A, Vanderschuren H, Thonar C & Delaplace P (2025) Trade-offs between agronomic yields and sustainability in winter wheat cropping systems under climate change mediated by soil organic matter content. PLoS Climate. 10.1371/journal.pclm.0000616.

11 Trnka, M., Dubrovský, M., Semerádová, D. et al. (2004). Projections of uncertainties in climate change scenarios into expected winter wheat yields. Theor Appl Climatol 77, 229–249 10.1007/s00704-004-0035-x.

12 Lal, R. (2020). Soil organic matter content and crop yield. Journal of Soil and Water Conservation, 75(2), 27A–32A. 10.2489/jswc.75.2.27A.

13 Marañón-Jiménez S, Luo X, Richter A, Gündler P, Fuchslueger L, Verbrigghe N, Poeplau C, Sigurdsson BD, Janssens I, Peñuelas J. (2025). Warming Weakens Soil Nitrogen Stabilization Pathways Driving Proportional Carbon Losses in Subarctic Ecosystems. Glob Chang Biol. 31:e70309. doi: 10.1111/gcb.70309.

14. Roy J, Rineau F, De Boeck HJ, Nijs I, Pütz T, Abiven S, et al. (2021). Ecotrons: Powerful and versatile ecosystem analysers for ecology, agronomy and environmental science. Glob. Change Biol., 27: 1387–1407. 10.1111/gcb.15471.

15 Keuskamp, J. A., Dingemans, B. J., Lehtinen, T., Sarneel, J. M., & Hefting, M. M. (2013). Tea Bag Index: a novel approach to collect uniform decomposition data across ecosystems. Methods in Ecology and Evolution, 4(11), 1070–1075. 10.1111/2041-210X.12097.

16 Dreywood, R. (1946). Qualitative test for carbohydrate material. Industrial & Engineering Chemistry Analytical Edition, 18(8), 499–499. 10.1021/i560156a015.

17 Campbell CD, Chapman SJ, Cameron CM, Davidson MS, Potts JM. (2002). A rapid microtiter plate method to measure carbon dioxide evolved from carbon substrate amendments so as to determine the physiological profiles of soil microbial communities by using whole soil. Appl Environ Microbiol (6):3593-9. 10.1128/AEM.69.6.3593-3599.2003.

18 Vance, E. D., Brookes, P. C., & Jenkinson, D. S. (1987). An extraction method for measuring soil microbial biomass C. Soil biology and Biochemistry, 19(6), 703–707. 10.1016/0038-0717(87)90052-6.

19 Fierer N, Jackson JA, Vilgalys R, Jackson RB. (2005). Assessment of soil microbial community structure by use of taxon-specific quantitative PCR assays. Appl Environ Microbiol (7):4117–20. 10.1128/AEM.71.7.4117-4120.2005.

20 Chemidlin Prévost-Bouré N, Christen R, Dequiedt S, Mougel C, Lelièvre M, Jolivet C, et al. (2011). Validation and Application of a PCR Primer Set to Quantify Fungal Communities in the Soil Environment by Real-Time Quantitative PCR. PLoS ONE 6(9): e24166. 10.1371/journal.pone.0024166.

21 Liu C.M., Aziz M., Kachur S., Hsueh P.-R., Huang Y.-T., Keim P. & Price L.B., 2012. BactQuant:An enhanced broad-coverage bacterial quantitative real-time PCR assay. BMC Microbiol 12(1), 56. 10.1186/1471-2180-12-56.

22 Carrascal, L.M., Galván, I. and Gordo, O. (2009), Partial least squares regression as an alternative to current regression methods used in ecology. Oikos, 118: 681–690. 10.1111/j.1600-0706.2008.16881.x.

23 Akselrud, C. I. A. (2024). Random forest regression models in ecology: Accounting for messy biological data and producing predictions with uncertainty. Fisheries Research, 280, 107161. 10.1016/j.fishres.2024.107161.

24. R Core Team (2025). R: A Language and Environment for Statistical Computing. R Foundation for Statistical Computing, Vienna, Austria. https://www.R-project.org.

25. Fox J, Weisberg S (2019). An R Companion to Applied Regression, Third edition. Sage, Thousand Oaks CA. https://www.john-fox.ca/Companion/.

26 Kuhn, M. (2008). Building Predictive Models in R Using the caret Package. Journal of Statistical Software, 28(5), 1–26. 10.18637/jss.v028.i05.

27 Wickham H, François R, Henry L, Müller K, Vaughan D (2023). dplyr: A Grammar of Data Manipulation. 10.32614/CRAN.package.dplyr.

28 Fox, J. (2003). Effect Displays in R for Generalised Linear Models. Journal of Statistical Software, 8(15), 1–27. 10.18637/jss.v008.i15.

29 Lenth R, Piaskowski J (2025). emmeans: Estimated Marginal Means, aka Least-Squares Means. 10.32614/CRAN.package.emmeans.

30 Campitelli E (2025). ggnewscale: Multiple Fill and Colour Scales in ’ggplot2’. 10.32614/CRAN.package.ggnewscale.

31. Wickham, H. (2016). ggplot2: Elegant Graphics for Data Analysis. Springer-Verlag New York, 2016. https://ggplot2.tidyverse.org.

32 Bates D, Maechler M, Bolker B, Walker S (2015). Fitting Linear Mixed-Effects Models Using lme4. Journal of Statistical Software, 67(1), 1–48. 10.18637/jss.v067.i01.

33 Kuznetsova A, Brockhoff PB, Christensen RHB (2017). lmerTest Package: Tests in Linear Mixed Effects Models. Journal of Statistical Software, 82(13), 1–26. 10.18637/jss.v082.i13.

34 Pedersen T (2025). patchwork: The Composer of Plots. 10.32614/CRAN.package.patchwork.

35 Lüdecke, et al. (2021). performance: An R Package for Assessment, Comparison and Testing of Statistical Models. Journal of Open Source Software, 6(60), 3139. 10.21105/joss.03139.

36 Liland K, Mevik B, Wehrens R (2024). pls: Partial Least Squares and Principal Component Regression. 10.32614/CRAN.package.pls.

37 Mehmood, T., Liland, K.H., Snipen L., Sæbø S (2012). A review of variable selection methods in Partial Least Squares Regression. Chemometrics and Intelligent Laboratory Systems 118, pp. 62–69.

38 Liaw A, Wiener M (2002). Classification and Regression by randomForest. R News, 2(3), 18–22. https://CRAN.R-project.org/doc/Rnews/.

39 Müller K, Wickham H (2025). tibble: Simple Data Frames. 10.32614/CRAN.package.tibble.

40 Wickham H, Vaughan D, Girlich M (2024). tidyr: Tidy Messy Data. 10.32614/CRAN.package.tidyr.

41 Wickham H, Averick M, Bryan J, Chang W, McGowan LD, François R, Grolemund G, Hayes A, Henry L, Hester J, Kuhn M, Pedersen TL, Miller E, Bache SM, Müller K, Ooms J, Robinson D, Seidel DP, Spinu V, Takahashi K, Vaughan D, Wilke C, Woo K, Yutani H (2019). Welcome to the tidyverse. Journal of Open Source Software, 4(43), 1686. 10.21105/joss.01686.

42. Barel J. M., Kuyper T. W., de Boer W., & De Deyn G. B. (2019). Plant presence reduces root and shoot litter decomposition rates of crops and wild relatives. Plant and Soil, 438(1), 313–327. 10.1007/s11104-019-03981-7.

43 Bever JD, Dickie IA, Facelli E, Facelli JM, Klironomos J, Moora M, Rillig MC, Stock WD, Tibbett M, Zobel M. (2010). Rooting theories of plant community ecology in microbial interactions. Trends Ecol Evol (8):468-78. 10.1016/j.tree.2010.05.004.

44 Chung, Y. A., Ke, P.-J., & Adler, P. B. (2023). Mechanistic approaches to investigate soil microbe-mediated plant competition. Journal of Ecology, 111, 1590–1597. 10.1111/1365-2745.14156.

45 Zhang M, Zhang L, Li J, Huang S, Wang S, Zhao Y, Zhou W, Ai C. (2025). Nitrogen-shaped microbiotas with nutrient competition accelerate early-stage residue decomposition in agricultural soils. Nat Commun. 2025 Jul 1;16(1):5793. 10.1038/s41467-025-60948-2.

46 Dijkstra, F. A., Bader, N. E., Johnson, D. W., & Cheng, W. (2009). Does accelerated soil organic matter decomposition in the presence of plants increase plant N availability?. Soil Biology and Biochemistry, 41(6), 1080–1087. 10.1016/j.soilbio.2009.02.013.

47 Michel, J., Fontaine, S., Revaillot, S., Piccon-Cochard, C., & Whitaker, J. (2024). Vegetative stage and soil horizon respectively determine direction and magnitude of rhizosphere priming effects in contrasting tree line soils. Functional Ecology, 38, 1984–2002. 10.1111/1365-2435.14625.

48 Lian, J., Massart, S., Li, G., & Zhang, J. (2026). Root processes counteract the suppression of nitrogen-induced priming effects by enhancing microbial activity and catabolism in greenhouse vegetable production systems. Soil and Tillage Research, 255, 106802. 10.1016/j.still.2025.106802.

49 Nacry, P., Bouguyon, E. & Gojon, A. (2013). Nitrogen acquisition by roots: physiological and developmental mechanisms ensuring plant adaptation to a fluctuating resource. Plant Soil 370, 1–29. 10.1007/s11104-013-1645-9.

50. Waibel M, Michel J, Antoine M, Balanzategui-Guijarro I, Cao D, Delaplace P, Manfroy S, Moya-Laraño J, Sanchez-Moreno S, Santin-Montanya I, Tenorio JL, Thonar C, Vanderschuren H, Van Der Straeten D, Verlinde T, Weinmann M & Symanczik S. (2025) Reduction in tillage intensity associates with lower wheat yields and grain nutrients despite improved soil biological indicators. Agriculture, Ecosystems & Environment 389, 109675. 10.1016/j.agee.2025.109675.

51. Moore J. C., McCann K., & de Ruiter P. C. (2005). Modeling trophic pathways, nutrient cycling, and dynamic stability in soils. Pedobiologia, 49(6), 499–510. 10.1016/j.pedobi.2005.05.008.

52 Hines J, Megonigal JP, Denno RF. (2006). Nutrient subsidies to belowground microbes impact aboveground food web interactions. Ecology 87(6):1542–55. 10.1890/0012-9658(2006)87[1542:nstbmi]2.0.co;2.

53 Potapov AM. (2022). Multifunctionality of belowground food webs: resource, size and spatial energy channels. Biol Rev Camb Philos Soc 97(4):1691–1711. 10.1111/brv.12857.

54 Shaposhnikov, A.I., Belimov, A.A., Azarova, T.S. et al. (2023). The Relationship Between the Composition of Root Exudates and the Efficiency of Interaction of Wheat Plants with Microorganisms. Appl Biochem Microbiol 59, 330–343. 10.1134/S000368382303016X.

55. Michel J, Balanzategui-Guijarro I, Cao D, Hinsinger P, Le Gouis J, Moya-Laraño J, Sánchez-Moreno S, Symanczik S, Vanderschuren H, Van Der Straeten D, Waibel M, Weinmann M, Thonar C & Delaplace P (2025) Sustainable and resilient agroecosystems need complexity of soil food webs and multivariate soil health indicators. European Journal of Soil Science 76, no. 5: e70192. 10.1111/ejss.70192.

56 Wu, Z., Liu, Z., Wang, W. et al. (2026). Root-knot nematode Meloidogyne incognita uses secondary-metabolite-mediated soil microbiome shifts to locate host plants. Nat. Plants 10.1038/s41477-025-02205-4.

57 Zhao, D., Qiu, J., Fan, Z., Ti, C., Huang, Z., Yan, X., & Xia, Y. (2025). Global warming has imbalance impact on soil nitrogen transformation rates. Earth’s Future, 13, e2024EF004756. 10.1029/2024EF004756.

58 Mao, C., Wang, Y., Ran, J., Wang, C., Yang, Z., & Yang, Y. (2025). Effects of warming and precipitation change on soil nitrogen cycles: a meta-analysis. Journal of Plant Ecology, 18(3), rtaf051. 10.1093/jpe/rtaf051.

59 Barłóg P, Grzebisz W, Łukowiak R. (2022). Fertilizers and Fertilization Strategies Mitigating Soil Factors Constraining Efficiency of Nitrogen in Plant Production. Plants (Basel). 11(14):1855. 10.3390/plants11141855.

60 Koller, R., Scheu, S., Bonkowski, M., Robin, C. (2013). Protozoa stimulate N uptake and growth of arbuscular mycorrhizal plants, Soil Biology and Biochemistry 65, 204–210. 10.1016/j.soilbio.2013.05.020.

61 Yu A. (2024). Rhizosphere Dynamics: Unraveling the Complex Interactions Beneath the Soil. International Research Journal of Plant Science Vol. 15(3), 1–2. DOI: 10.14303/irjps.2024.25.

62 Siddique, M. B. A., Khalid, A., Ditta, A., Mahmood, S., Alataway, A., Dewidar, A. Z., & Mattar, M. A. (2023). Climate change variables modify microbial community structure and soil enzymes involved in nitrogen and phosphorus metabolism. Rhizosphere, 28, 100793. 10.1016/j.rhisph.2023.100793.

63 Liu S, Ru J, Guo X, Gao Q, Deng S, Lei J, Song J, Zhai C, Wan S, Yang Y. (2025). Altered precipitation and nighttime warming reshape the vertical distribution of soil microbial communities. mSystems 10:e01248–24. 10.1128/msystems.01248-24.

64 Zhang K, Shi Y, Jing X, He JS, Sun R, Yang Y, Shade A, Chu H. (2026). Effects of Short-Term Warming and Altered Precipitation on Soil Microbial Communities in Alpine Grassland of the Tibetan Plateau. Front Microbiol. 7:1032. doi: 10.3389/fmicb.2016.01032. Erratum in: Front Microbiol. 2017 Apr 18;8:667. doi: 10.3389/fmicb.2017.00667. 10.3389/fmicb.2016.01032.

65 Knight, C.G., Nicolitch, O., Griffiths, R.I., Goodall, T., Jones, B., Weser, C. et al. (2024). Soil microbiomes show consistent and predictable responses to extreme events. Nature 636, 690–696 (2024). 10.1038/s41586-024-08185-3.

66 Drake, J. E., Darby, B. A., Giasson, M.-A., Kramer, M. A., Phillips, R. P., & Finzi, A. C. (2013). Stoichiometry constrains microbial response to root exudation- insights from a model and a field experiment in a temperate forest, Biogeosciences, 10, 821–838, 10.5194/bg-10-821-2013.

67 Wei T, Zhang P, Wang K, Ding R, Yang B, Nie J, et al. (2015). Effects of Wheat Straw Incorporation on the Availability of Soil Nutrients and Enzyme Activities in Semiarid Areas. PLoS ONE 10(4): e0120994. 10.1371/journal.pone.0120994.

68 Weil, R.R. and Brady, N.C. (2017) The Nature and Properties of Soils. 15th Edition, Pearson, New York. ISBN 13: 978-1-292-16223-2.

69 Middelanis, T., Pohl, C. M., Looschelders, D., & Hamer, U. (2023). New directions for the Tea Bag Index: Alternative teabags and concepts can advance citizen science. Ecological Research, 38(5), 690–699. 10.1111/1440-1703.12409.

70 Marschner, P., Umar, S., & Baumann, K. (2011). The microbial community composition changes rapidly in the early stages of decomposition of wheat residue. Soil Biology and Biochemistry, 43(2), 445–451. 10.1016/j.soilbio.2010.11.015.

71 Wahdan SFM, Hossen S, Tanunchai B, Sansupa C, Schädler M, Noll M, Dawoud TM, Wu YT, Buscot F, Purahong W. Life in the Wheat Litter: Effects of Future Climate on Microbiome and Function During the Early Phase of Decomposition. Microb Ecol. 2022 Jul;84(1):90–105. 10.1007/s00248-021-01840-6.

72. Geisen S., Wall D.H., & van der Putten W.H. (2018). Soil protists: a fertile frontier in soil biology research. FEMS Microbiology Reviews, 42(3), 293–323. 10.1093/femsre/fuy006.

73 Mielke, L., Taubert, M., Cesarz, S., Ruess, L., Kuesel, K., Gleixner, G., & Lange, M. (2022). Nematode grazing increases the allocation of plant-derived carbon to soil bacteria and saprophytic fungi, and activates bacterial species of the rhizosphere. Pedobiologia, 90, 150787. 10.1016/j.pedobi.2021.150787.

74 Henriksen, T. M., & Breland, T. A. (1999). Nitrogen availability effects on carbon mineralization, fungal and bacterial growth, and enzyme activities during decomposition of wheat straw in soil. Soil Biology and Biochemistry, 31(8), 1121–1134. 10.1016/S0038-0717(99)00030-9.

75 Sierra, C. A., Malghani, S., and Loescher, H. W. (2017). Interactions among temperature, moisture, and oxygen concentrations in controlling decomposition rates in a boreal forest soil, Biogeosciences, 14, 703–710, 10.5194/bg-14-703-2017.

76 Evans, S. E., Allison, S. D., & Hawkes, C. V. (2022). Microbes, memory and moisture: Predicting microbial moisture responses and their impact on carbon cycling. Functional Ecology, 36, 1430–1441. 10.1111/1365-2435.14034.

77 Qin S. et al. (2019) Temperature sensitivity of SOM decomposition governed by aggregate protection and microbial communities.Sci.Adv.5,eaau1218(2019). 10.1126/sciadv.aau1218.

78 Moinet, G. Y., Hunt, J. E., Kirschbaum, M. U., Morcom, C. P., Midwood, A. J., & Millard, P. (2018). The temperature sensitivity of soil organic matter decomposition is constrained by microbial access to substrates. Soil Biology and Biochemistry, 116, 333–339. 10.1016/j.soilbio.2017.10.031.

79 Carey JC, Tang J, Templer PH, Kroeger KD, Crowther TW, Burton AJ, et al. (2016). Temperature response of soil respiration largely unaltered with experimental warming. Proc Natl Acad Sci U S A. 2016 Nov 29;113(48):13797-13802. 10.1073/pnas.1605365113.

80 Robinson SI, O’Gorman EJ, Frey B, Hagner M, Mikola J. Soil organic matter, rather than temperature, determines the structure and functioning of subarctic decomposer communities. Glob Chang Biol. 2022 Jun;28(12):3929–3943. 10.1111/gcb.16158.

81 Liu F, Zhang W, Li S. (2025). Effects of Freeze-Thaw Cycles on Uptake Preferences of Plants for Nutrient: A Review. Plants 14(7):1122. 10.3390/plants14071122.

82 Yang, YF., Wang, CX., He, XL. et al. (2025). Study on the effects of winter irrigation during seasonal freezing–thawing period on soil microbial ecological properties. Sci Rep 15, 23586. 10.1038/s41598-025-07845-2.

83 Shi Y, Lalande R, Hamel C, Ziadi N. (2015). Winter effect on soil microorganisms under different tillage and phosphorus management practices in eastern Canada. Can J Microbiol. 61(5):315–26. 10.1139/cjm-2014-0821.

84 Han, J., Xing, L., Zhang, C., Li, J., Li, Y., Zhang, Y., He, H., Hu, C., Li, X., Zhang, L., Dong, W., Qin, S., & Liu, X. (2024). The Effect of Soil Microbial Residues-Mediated Nitrogen Conservation and Supply during the Growing Season on Nitrogen Uptake by Wheat. Agronomy, 14(1), 193. 10.3390/agronomy14010193.

85 Volpe, V., Marani, M., Albertson, J. D., & Katul, G. (2013). Root controls on water redistribution and carbon uptake in the soil–plant system under current and future climate. Advances in Water resources, 60, 110–120. 10.1016/j.advwatres.2013.07.008.

86 Verhoef, A., & Egea, G. (2014). Modeling plant transpiration under limited soil water: Comparison of different plant and soil hydraulic parameterizations and preliminary implications for their use in land surface models. Agricultural and Forest Meteorology, 191, 22–32. 10.1016/j.agrformet.2014.02.009.

87 Feifel, M., Durner, W., Hohenbrink, T. L., & Peters, A. (2024). Effects of improved water retention by increased soil organic matter on the water balance of arable soils: A numerical analysis. Vadose Zone Journal, 23, e20302. 10.1002/vzj2.20302.

88 Oldfield, E. E., Bradford, M. A., & Wood, S. A. (2019). Global meta-analysis of the relationship between soil organic matter and crop yields. Soil, 5(1), 15–32. 10.5194/soil-5-15-2019.

89 Xu X, Hui D, King AW, Song X, Thornton PE, Zhang L. Convergence of microbial assimilations of soil carbon, nitrogen, phosphorus, and sulfur in terrestrial ecosystems. Sci Rep. 2015 Nov 27;5:17445. 10.1038/srep17445.

90 Su, B., & Shangguan, Z. (2022). Stoichiometric homeostasis in response to variable water and nutrient supply in a Robinia pseudoacacia plant–soil system. Journal of Plant Ecology, 15(5), 991–1006. 10.1093/jpe/rtac011.

91. Cao D, Michel J, Lorer E, Weinmann M, Le Gouis J, Leon C, Perrochon S, Alvarez D, Leemans V, Balanzategui I, Moya Laraño J, Sanchez Moreno S, Symanczik S, Waibel M, Vanderschuren H, Thonar C, Delaplace P & Van Der Straeten D (2026) Climate change threatens nutritional quality of European winter wheat. Preprint under review.

92 Yousefi, M., Dray, A., & Ghazoul, J. (2024). Assessing the effectiveness of cover crops on ecosystem services: a review of the benefits, challenges, and trade-offs. International Journal of Agricultural Sustainability, 22(1). 10.1080/14735903.2024.2335106.

93 Averill, C., Turner, B. & Finzi, A. (2014). Mycorrhiza-mediated competition between plants and decomposers drives soil carbon storage. Nature 505, 543–545. 10.1038/nature12901.

94 Averill, C. (2016). Slowed decomposition in ectomycorrhizal ecosystems is independent of plant chemistry. Soil Biology and Biochemistry, 102, 52–54. 10.1016/j.soilbio.2016.08.003.

95. Peng, Y., Rieke, E. L., Chahal, I., Norris, C. E., Janovicek, K., Mitchell, J. P., … & Van Eerd, L. L. (2023). Maximizing soil organic carbon stocks under cover cropping: Insights from long-term agricultural experiments in North America. Agriculture, Ecosystems & Environment, 356, 108599. 10.1016/j.agee.2023.108599.

96 Zhu, S., Sainju, U. M., Zhang, S., Tan, G., Wen, M., Dou, Y., … & Wang, J. (2024). Cover cropping promotes soil carbon sequestration by enhancing microaggregate-protected and mineral-associated carbon. Science of the Total Environment, 908, 168330. 10.1016/j.scitotenv.2023.168330.

97 Bozal-Leorri, A., Corrochano-Monsalve, M., Arregui, L.M. et al. (2021). Biological and synthetic approaches to inhibiting nitrification in non-tilled Mediterranean soils. Chem. Biol. Technol. Agric. 8, 51. 10.1186/s40538-021-00250-7.

98 Lan, T., Li, M., He, X. et al. (2022). Effects of synthetic nitrification inhibitor (3,4-dimethylpyrazole phosphate; DMPP) and biological nitrification inhibitor (methyl 3-(4-hydroxyphenyl) propionate; MHPP) on the gross N nitrification rate and ammonia oxidizers in two contrasting soils. Biol Fertil Soils 58, 333–344. 10.1007/s00374-022-01628-x.

99 Tajima, H., Yadav, A., Castellanos, J.H., Yan, D., Brookbank, B.P., Nambara, E. and Blumwald, E. (2025). Increased Apigenin in DNA-Edited Hexaploid Wheat Promoted Soil Bacterial Nitrogen Fixation and Improved Grain Yield Under Limiting Nitrogen Fertiliser. Plant Biotechnol. J, 23: 5146–5160. 10.1111/pbi.70289.

100 Radicetti, E., Baresel, J. P., El-Haddoury, E. J., Finckh, M. R., Mancinelli, R., Schmidt, J. H., … & Campiglia, E. (2018). Wheat performance with subclover living mulch in different agro-environmental conditions depends on crop management. European Journal of Agronomy, 94, 36–45. 10.1016/j.eja.2018.01.011.

101 Fontaine, D., Eriksen, J., & Sørensen, P. (2020). Cover crop and cereal straw management influence the residual nitrogen effect. European Journal of Agronomy, 118, 126100. 10.1016/j.eja.2020.126100.

102 Cao, Y., Yin, T., Zhang, Y., Yang, X., Liu, B., Zhu, Y., … & Liu, L. (2024). Quantitative assessment of the effects of rising temperature on the grain protein of winter wheat in china and its adaptive strategies. Computers and Electronics in Agriculture, 226, 109474. 10.1016/j.compag.2024.109474.

103 Cheng, W., Johnson, D.W. and Fu, S. (2003), Rhizosphere Effects on Decomposition. Soil Sci. Soc. Am. J., 67: 1418–1427. 10.2136/sssaj2003.1418.

