## Supplementary meterial for tea time for "Teatime for *Triticum* – (how) can the presence of plants slow down decomposition?"

**Supplementary material for teatime**

SM0: Tea bag deployment scheme

SM1: Soil moisture (%) in planted and unplanted soils at DAS131 & DAS 159

SM2: Details of qPCR

SM3: Parameters included in PLS-VIP

SM4: Plot of top ranked VIPs

SM5: Parameter scores in RF model

SM6: RF variable importance

SM7: Partial dependence plots

SM8: Decomposition of green and rooibos tea

**SM0: Tea bag deployment scheme**

|  |  | 1 |  | 2 |  | 3 |
| --- | --- | --- | --- | --- | --- | --- |
|  |  |  | 4 |  | 5 |  |
|  |  | 6 |  | 7 |  | 8 |
|  |  |  | 9 |  | 10 |  |
|  |  | door | | | | |
|  |  | depth = 8cm | | |  |  |
|  |  | unplanted installed 02/05 (131 DAS) | | | | |
|  |  | planted installed 03/05 (132 DAS) | | | | |

**SM1: Soil moisture (%) in planted and unplanted soils at DAS131 & DAS 159**

| Humidity131 | Hum159 | Av.hum | Soil | Plant | Climate | 131.delta | 159.delta | Av.delta |
| --- | --- | --- | --- | --- | --- | --- | --- | --- |
| 20.50 | 17.32 | 18.91 | S1 | Yes | 2013 | 2.84 | 2.64 | 2.74 |
| 23.34 | 19.96 | 21.65 | S1 | No | 2013 |  |  |  |
| 17.25 | 18.42 | 17.84 | S1 | Yes | 2068 | 3.47 | NA | 2.88 |
| 20.72 | NA | 20.72 | S1 | No | 2068 |  |  |  |
| 15.69 | 9.94 | 12.81 | S1 | Yes | 2085 | 2.56 | 5.30 | 3.93 |
| 18.25 | 15.23 | 16.74 | S1 | No | 2085 |  |  |  |
| 23.14 | 20.70 | 21.92 | S2 | Yes | 2013 | -11.58 | -11.59 | -11.58 |
| 11.56 | 9.11 | 10.34 | S2 | No | 2013 |  |  |  |
| 19.97 | 19.57 | 19.77 | S2 | Yes | 2068 | -3.44 | -0.41 | -1.92 |
| 16.53 | 19.17 | 17.85 | S2 | No | 2068 |  |  |  |
| 19.55 | 11.14 | 15.35 | S2 | Yes | 2085 | 2.21 | 6.37 | 4.29 |
| 21.76 | 17.51 | 19.64 | S2 | No | 2085 |  |  |  |

**SM2: Details of qPCR**

Quantitative PCR (qPCR) was conducted on sieved soil samples. Approximately 2g of each soil sample was freeze-dried after sieving and stored at -20°C. DNA extraction was performed using the DNeasy PowerLyzer PowerSoil Kit (QIAGEN, Germany) and its protocol. The extracted DNA was quantified using a Quantus™ quantifluorimeter (Promega Corporation, United States). A stock solution was prepared by diluting the 20X buffer with DNA-free water, and the DNA sample was added to this solution for fluorescence reading. The sample DNA was diluted to a final concentration of 10 ng mL^-1^. Thereafter, a qPCR reaction mix was prepared by combining 5 μl of SYBR Green with two specific primers (1 μl each) and with 2 μl of sterile water devoid of DNA. SYBR Green is a fluorescent dye that specifically interacts with double-stranded DNA. When present in the reaction mixture, it binds to DNA amplified during qPCR and emits fluorescence proportional to the amount of DNA present in the reaction. In this case, the employed primers included the rRNA 16s primer, BactQuant (Liu et al., 2012), for bacterial quantification, and the rRNA 18s primer, FR1/FF390 (Chemidlin Prévost-Bouré et al., 2011), for quantifying fungal presence in the samples. Subsequently, this reaction mixture was precisely deposited at the bottom of the wells of the qPCR MicroAmp™ Optical 96-Well Reaction Plates (Thermo Fisher Scientific Inc., United States), where 1 μl of the DNA sample, having a target concentration of 10 ng/mL, was deposited on the edges of the wells containing the solution. To avoid the risk of contamination and evaporation, a protective film (Applied Biosystems™ MicroAmp™ Optical Adhesive Film, Thermo Fisher Scientific Inc., United States) was carefully applied to the plate. The prepared plates were then placed in a QuantiStudio 3 thermocycler (Thermo Fisher Scientific Inc., United States) to perform a series of 40 cycles of qPCR. Each cycle includes denaturation, hybridisation and extension steps, which allow specific amplification of the target region of DNA. During this process, the fluorescence emitted by the SYBR Green was measured at each cycle to follow in real time the amplification of the DNA and to allow the quantitative analysis of the results. A calibration curve was established by proceeding in the same way as above, except that the DNA was substituted by a series of dilutions of a plasmid containing the DNA sequence of interest to be quantified. The data was then analysed using the Applied QuantiStudio 3 software (Thermo Fisher Scientific Inc., United States).

**SM3: Parameter scores in PLS-VIP**

| Loadings: | Comp 1 | Comp 2 | Comp 3 | Comp 4 | Comp 5 |
| --- | --- | --- | --- | --- | --- |
| Number.of.Root.Tips |  | -0.178 | 0.399 | -0.267 | -0.163 |
| Number.of.Branch.Points |  | -0.177 | 0.404 | -0.237 | -0.169 |
| Total.Root.Length.mm |  | -0.173 | 0.4 | -0.272 | -0.15 |
| Branching.frequency.per.mm |  | -0.261 | 0.336 | -0.236 |  |
| Network.Area.mm2 |  | -0.177 | 0.391 | -0.293 | -0.139 |
| CO2.basal.102 | -0.258 | 0.322 | -0.121 |  |  |
| CO2.exudate.102 | -0.22 | 0.113 | -0.236 |  | -0.161 |
| CO2.basal.131 | -0.108 | 0.282 | -0.122 |  | -0.239 |
| CO2.exudate.131 |  | 0.236 | -0.143 |  | -0.267 |
| CO2.basal.159 | -0.293 |  |  |  | -0.151 |
| CO2.exudate.159 | -0.276 |  |  | 0.159 | -0.172 |
| NO3.mg.L.82 | -0.152 | 0.31 |  | -0.227 | 0.318 |
| Glu.mg.L.82 | -0.166 |  |  | -0.358 | 0.383 |
| NO3.mg.L.102 |  | 0.261 | -0.114 |  |  |
| Glu.mg.L.102 |  |  | 0.28 | 0.294 | -0.315 |
| NO3.mg.L.110 | -0.27 | 0.167 | 0.123 | -0.263 | 0.178 |
| Glu.mg.L.110 | -0.168 | 0.163 | 0.118 | -0.142 | 0.358 |
| NO3.mg.L.131 | -0.147 | 0.335 |  | -0.239 | 0.182 |
| Glu.mg.L.131 |  |  |  | 0.13 | 0.165 |
| NO3.mg.L.150 | -0.104 | 0.332 |  |  |  |
| Glu.mg.L.150 | -0.145 | 0.104 | 0.314 | -0.125 | 0.188 |
| mbC.mig.g.1.131 | -0.228 | 0.155 | 0.231 | -0.232 | 0.126 |
| bacteria_gene.copies.g.1.131 | 0.321 |  |  | -0.207 | 0.219 |
| fungi_gene.copies.g.1.131 | 0.392 |  |  | -0.105 | -0.124 |
| FB.131 |  | -0.274 |  |  |  |
| mbC.mig.g.1.159 |  | 0.177 |  |  | -0.195 |
| bacteria_gene.copies.g.1.159 | 0.109 |  |  | -0.138 | -0.269 |
| fungi_gene.copies.g.1.159 | 0.315 |  |  | -0.415 |  |
| FB.159 | 0.322 |  |  |  | 0.197 |
| delta.mbC | 0.118 |  | -0.175 | 0.219 | -0.244 |
| SOCg.kg |  | 0.145 | -0.373 |  | 0.127 |
| Sand | -0.193 | 0.153 |  |  | 0.102 |
| Cum.rain |  | -0.279 | 0.105 |  | 0.378 |
| Av.soil.MC | -0.291 |  |  |  | 0.221 |
| Av.soilT |  | 0.321 | -0.225 |  |  |
| Av.airT | -0.179 | 0.213 |  |  | -0.462 |
| SS loadings | 1.23 | 1.333 | 1.431 | 1.204 | 1.667 |
| Proportion Var | 0.034 | 0.037 | 0.04 | 0.033 | 0.046 |
| Cumulative Var | 0.034 | 0.071 | 0.111 | 0.144 | 0.191 |

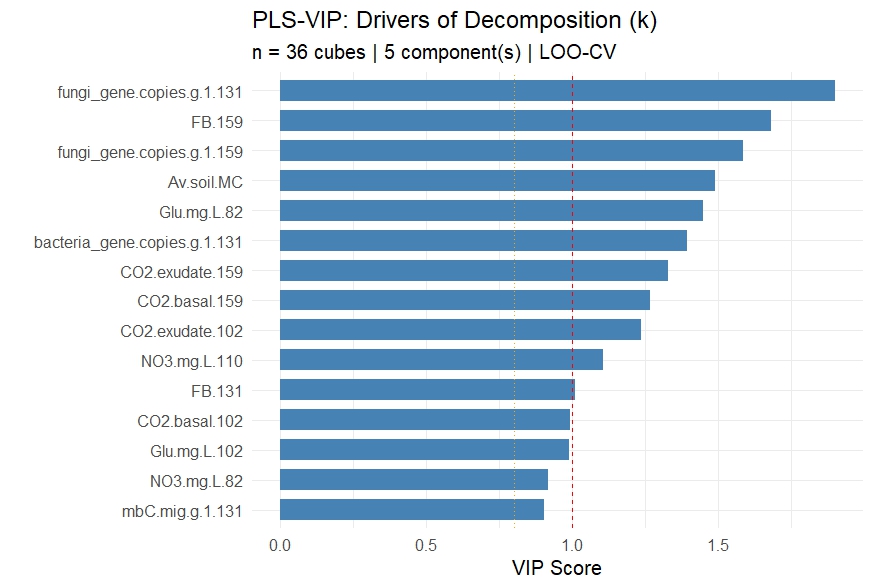
**SM4: Plot of top ranked VIP’s**

**SM5: Parameter scores in RF model**

|  | %IncMSE | IncNodePurity |
| --- | --- | --- |
| Number.of.Root.Tips | 4.066144 | 2.81E-05 |
| Number.of.Branch.Points | 3.008219 | 1.63E-05 |
| Total.Root.Length.mm | 4.103017 | 3.12E-05 |
| Branching.frequency.per.mm | 3.311495 | 2.07E-05 |
| Network.Area.mm2 | 3.901773 | 2.55E-05 |
| CO2.basal.102 | 7.543773 | 1.74E-04 |
| CO2.exudate.102 | 11.10383 | 1.14E-03 |
| CO2.basal.131 | 5.400498 | 1.85E-04 |
| CO2.exudate.131 | 6.133002 | 1.84E-04 |
| CO2.basal.159 | 10.21627 | 9.82E-04 |
| CO2.exudate.159 | 3.406716 | 9.86E-05 |
| NO3.mg.L.82 | 6.208499 | 5.87E-05 |
| Glu.mg.L.82 | 5.595478 | 2.84E-04 |
| NO3.mg.L.102 | 5.467547 | 3.52E-05 |
| Glu.mg.L.102 | 4.300974 | 6.03E-05 |
| NO3.mg.L.110 | 7.488511 | 4.12E-04 |
| Glu.mg.L.110 | 5.655789 | 5.09E-05 |
| NO3.mg.L.131 | 11.78807 | 1.56E-04 |
| Glu.mg.L.131 | 4.001373 | 4.20E-05 |
| NO3.mg.L.150 | 4.715577 | 6.53E-05 |
| Glu.mg.L.150 | 9.010376 | 7.92E-05 |
| mbC.mig.g.1.131 | 3.882688 | 5.50E-05 |
| bacteria_gene.copies.g.1.131 | 5.248368 | 1.77E-04 |
| fungi_gene.copies.g.1.131 | 18.06554 | 2.38E-03 |
| FB.131 | 3.719592 | 2.99E-05 |
| mbC.mig.g.1.159 | 11.55046 | 4.55E-04 |
| bacteria_gene.copies.g.1.159 | 7.581933 | 2.72E-04 |
| fungi_gene.copies.g.1.159 | 6.153914 | 1.70E-04 |
| FB.159 | 5.839044 | 1.70E-04 |
| delta.mbC | 8.929542 | 3.03E-04 |
| SOCg.kg | 3.722823 | 2.39E-05 |
| Sand | 3.548828 | 7.22E-06 |
| Cum.rain | 3.500008 | 3.47E-05 |
| Av.soil.MC | 5.201149 | 2.62E-04 |
| Av.soilT | 3.344957 | 4.74E-05 |
| Av.airT | 3.142776 | 2.81E-05 |
| Final OOB MSE: 3.875828e-06 |  |  |
| Final OOB % Var explained: 0.9479193 | |  |

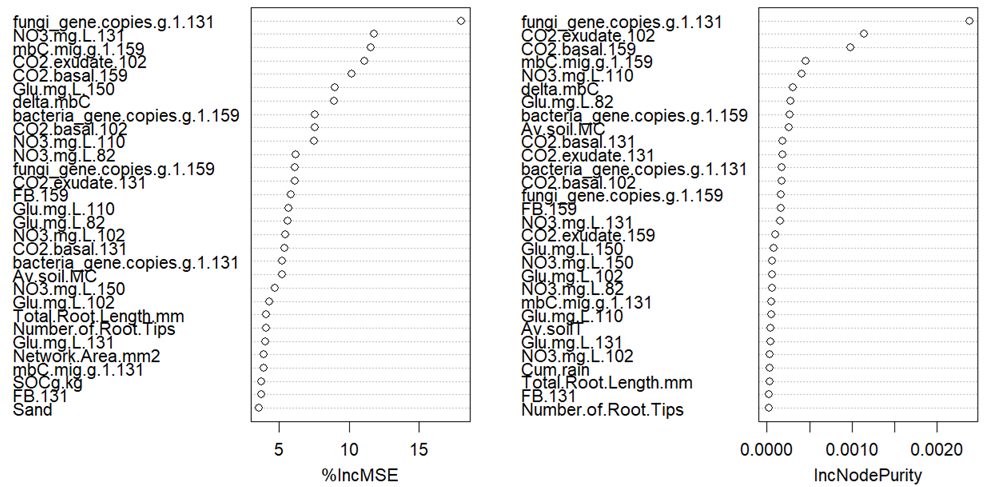
**SM6: RF variable importance**

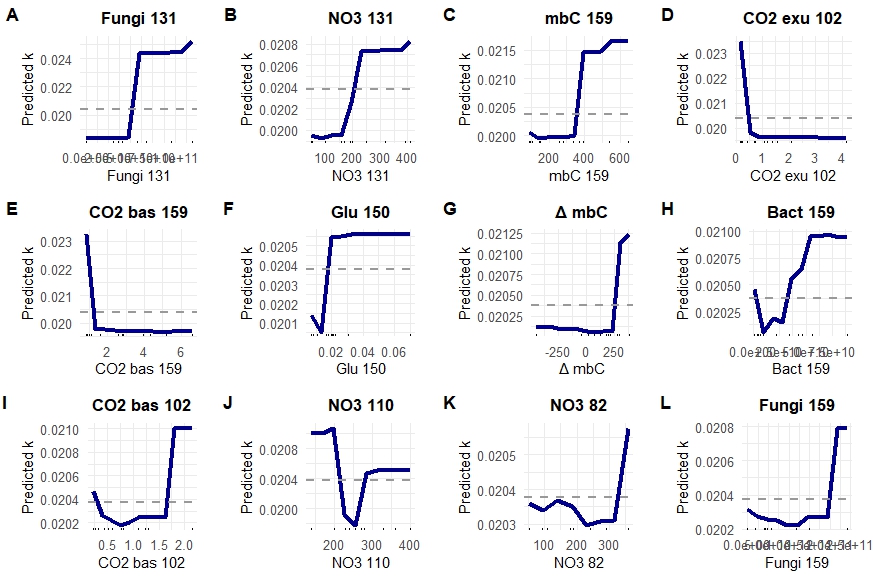
**SM7: Partial dependence plots**

**SM8: Decomposition of green and rooibos tea** (tea type p= 0.0001)

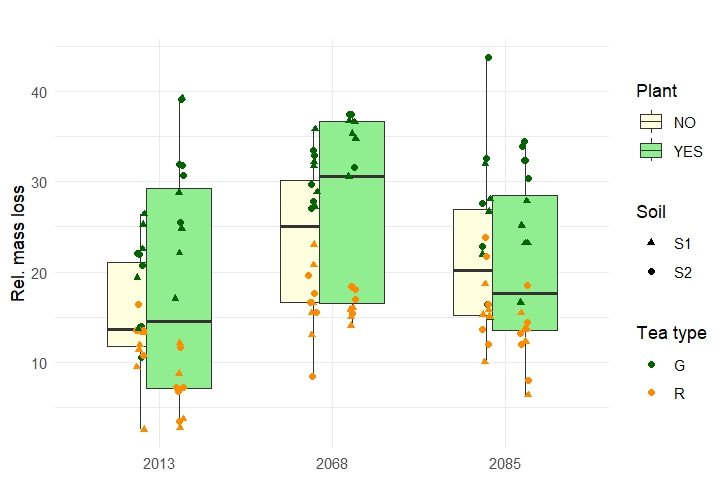
